# Convolutional Neural Networks for Decoding of Covert Attention Focus and Saliency Maps for EEG Feature Visualization

**DOI:** 10.1101/614784

**Authors:** Amr Farahat, Christoph Reichert, Catherine M. Sweeney-Reed, Hermann Hinrichs

## Abstract

**Objective:** Convolutional neural networks (CNNs) have proven successful as function approximators and have therefore been used for classification problems including electroencephalography (EEG) signal decoding for brain-computer interfaces (BCI). Artificial neural networks, however, are considered black boxes, because they usually have thousands of parameters, making interpretation of their internal processes challenging. Here we systematically evaluate the use of CNNs for EEG signal decoding and investigate a method for visualizing the CNN model decision process.

**Approach:** We developed a CNN model to decode the covert focus of attention from EEG event-related potentials during object selection. We compared the CNN and the commonly used linear discriminant analysis (LDA) classifier performance, applied to datasets with different dimensionality, and analyzed transfer learning capacity. Moreover, we validated the impact of single model components by systematically altering the model. Furthermore, we investigated the use of saliency maps as a tool for visualizing the spatial and temporal features driving the model output.

**Main results:** The CNN model and the LDA classifier achieved comparable accuracy on the lower-dimensional dataset, but CNN exceeded LDA performance significantly on the higher-dimensional dataset (without hypothesis-driven preprocessing), achieving an average decoding accuracy of 90.7% (chance level = 8.3%). Parallel convolutions, tanh or ELU activation functions, and dropout regularization proved valuable for model performance, whereas the sequential convolutions, ReLU activation function, and batch normalization components, reduced accuracy or yielded no significant difference. Saliency maps revealed meaningful features, displaying the typical spatial distribution and latency of the P300 component expected during this task.

**Significance:** Following systematic evaluation, we provide recommendations for when and how to use CNN models in EEG decoding. Moreover, we propose a new approach for investigating the neural correlates of a cognitive task by training CNN models on raw high-dimensional EEG data and utilizing saliency maps for relevant feature extraction.

## INTRODUCTION

Brain-computer interfaces (BCI) represent a bridge that allows direct communication between the brain and the environment without the need for muscular activity. They could be especially beneficial for patients who have lost muscular control and are ‘locked-in’, such as due to stroke or amyotrophic lateral sclerosis, or have sustained spinal cord injuries. To design such an interface, a voluntarily evoked neural signal must be recorded from the brain, and a pattern recognition system is required that can detect the signal.

Stimulus-evoked event-related potentials (ERP), such as the P300, are frequently elicited to provide the neural signal (40). The P300 is an attention-dependent ERP component showing a positive deflection in the ERP waveform, peaking 300 milliseconds after stimulus onset, regardless of the stimulus modality, e.g., visual, auditory, or somatosensory (38). It can be reliably evoked using the oddball paradigm, in which infrequent target “relevant” stimuli are presented among frequent standard “irrelevant” stimuli and is mainly distributed over the midline scalp electrodes (Fz, Cz, Pz), extending to parietal electrodes. This phenomenon was utilized to design the first P300 BCI that allowed participants to type letters by decoding the EEG signals and it was called “the P300 speller” (14). Other ERPs, such as visually evoked potentials (VEPs), can also be modulated by attention (13, 33). Both the P1 and the N1 components show larger amplitude for the attended stimuli than non-attended stimuli. The N1 component is followed by the P2 and N2 components that are elicited by rare, relevant stimuli (32). These modulations of VEPs depend on whether the participants are attending to the target stimulus overtly or covertly, i.e., whether they foveate the target stimulus or not. The classification performance and ERP analysis of P300 spellers under overt and covert attentional conditions were investigated by Treder and Blankertz (49), who concluded that P1, N1, P2, N2, and P3 components were enhanced in the case of overt attention. On the contrary, in the case of covert attention, only N2 and P3 were enhanced. As a result, the classifiers depend mainly on early VEPs in the case of overt attention and on late endogenous components in the case of covert attention. Moreover, it was shown that using posterior occipital electrodes Oz, PO7, and PO8 in addition to the regular frontal, central, and parietal electodes Fz, Cz, and Pz leads to significantly better classification performance of the P300 spellers (27). Comparing a full set of EEG electrodes to these customized electrodes showed that this significant increase in performance only occurs in the case of overt attention and not in the case of covert attention (6).

Machine learning techniques are effective in learning discriminative signal patterns from brain data. Most conventional machine learning techniques, however, require an initial step before classification, namely feature extraction. Feature extraction includes preprocessing like noise removal, dimensionality reduction and transformations like principal component analysis (PCA) or frequency decomposition, and feature selection like channel and time interval selection (2, 50, 51). Feature extraction is hypothesis-driven and requires domain expertise and thus using a machine learning algorithm that autonomously determines relevant features could save time and effort and potentially make use of features that would have been overlooked or removed.

Deep learning - a subarea of machine learning - has emerged in recent years as a technique that could be applied to extract features automatically from the data that maximize between-class discriminability (30). Deep learning has outperformed other machine learning techniques, especially on image and audio data, in a variety of tasks, by large margins (25, 42). Most of these successes are attributed to a specific type of deep learning models called convolutional neural networks (CNNs) (29). CNN refers to a special neural network architecture, which is inspired by the hierarchical structure of the ventral stream of the visual system (20). Stacking neurons with local receptive fields on top of each other leads to the development of simple and complex cells that extract features at different levels of abstraction (29).

The first attempt to use CNNs for P300 classification involved developing a 5-layer CNN (8), which was trained and tested on public dataset II of the third BCI competition (5). The method achieved the best results after 10 stimulus repetitions but the performance was not as high as that of an ensemble of support vector machines (SVMs) after 15 repetitions. The advantage of the CNN was that all available electrodes were used, instead of a priori electrode selection. A general CNN, EEGNet, was developed that can classify EEG signals for different BCI tasks (28). It follows the all-convolutional approach without fully connected layers before the output layer (45). It was trained and tested on a P300 dataset and yielded a significant improvement over classification using a shallow approach. Furthermore, a 3D CNN was developed to decode the P300 component in an auditory BCI (7). A 3D CNN is characterized by requiring the input as a 3D array: 2-dimensions for space and one time dimension. Instead of keeping the values of all electrodes at one time point in a single, one-dimensional array, the spatial structure of the data is further preserved. Comparison of the performance to that of a 2D version of the network and also to SVM and Fisher’s discriminant analysis (FDA) revealed significant improvement in decoding accuracy using the 3D CNN. Batch normalization is an algorithm that was developed to help facilitate faster training of neural networks with convergence on lower minima (22). A CNN was developed to examine the impact of including batch normalization on P300 detection accuracy (31). It was shown that removing more than one batch normalization layer markedly decreased the classification accuracy. CNNs trained on EEG data showed also a promising potential in decoding motor imagery tasks relative to the commonly used filter bank common spatial patterns (FBCSP) algorithm (28, 43).

In order to make CNNs interpretable, several feature visualization methods have been developed to explain the performance of the CNN models and the relevant features that drive their outputs (3, 35, 39). Some of these techniques were adopted and other novel techniques were developed for CNNs applied to EEG data (28, 43, 47). Here we adopt the gradient-based visualization technique commonly used in computer vision research called saliency maps (44). A saliency map is generated as the gradient of the CNN output with respect to its input example. It is computed using back propagation, after fixing the weights of the trained network, by propagating the gradient with respect to the layers’ inputs back to the first layer that receives the input data. We can interpret the saliency map representing the relevant features that drive the network’s decision.

Here we investigated the potential of a CNN model in decoding the focus of covert attention using EEG data, systematically tested the impact of different components of the model on its performance, and examined the model’s capacity for transfer learning. We explored the use of saliency maps for visualizing what the CNN model has learned from the data and investigated the possibility of using the features extracted by the CNN to gain insights into the electrophysiological signals underlying the task. Moreover, We compared the decoding performance of the CNN model to those obtained with a linear discriminant analysis (LDA) classifier, which is reported to be one of the best performing classifiers for ERP BCI applications (4, 26, 34), and also to the EEGNet model (28) to investigate the potential of CNNs in general in decoding the focus of covert attention.

## METHODS

### DATASET

The EEG data classified in the current study were recorded in a previous study, combined with simultaneous MEG recordings to compare both modalities for BCI use (41). In brief, nineteen participants (10 females) performed a covert attention task in multiple runs (10.7 runs per participant on average), each comprising 12 trials. In each trial, the participants covertly attended to one of 12 stimuli of their choice in an unpredictable order. The stimuli were presented as colored and numbered spherical objects at equidistant spaces over the circumference of an invisible circle with the fixation point located at its center.

All the objects flashed five times in a pseudorandom order, leading to 60 flashes with a stimulus onset asynchrony (SOA) of 167 ms. The pseudo-random order ensured that the same object would not flash twice in a row, but with at least two other objects flashing in between. The EEG data were recorded at 29 standard electrode locations (Fp1, Fp2, Fz, F3, F4, F7, F8, FC1, FC2, Cz, C3, C4, T7, T8, CP1, CP3, Pz,P3, P4, P7, P8, PO3, PO4, PO7, PO8, Oz, O9, O10, Iz), sampled at 508.63 Hz, and referenced against the right mastoid. Vertical and horizontal electrooculograms (EOG) were also recorded to track eye movements to ensure fixation of the gaze.

### PREPROCESSING

In order to test the potential of CNNs in dealing with high-dimensional EEG data, we created two datasets: one to represent high-dimensional, minimally preprocessed data and one to represent low-dimensional data preprocessed with a feature extraction method. For the first dataset, the data were only downsampled to half of the original sampling frequency (254.3 Hz) after applying an antialiasing FIR lowpass filter, and we call it dataset *f250*. For the second dataset, the data were decimated by a factor of 10 by averaging over blocks of 10 time points, which essentially is equivalent to downsampling after applying a moving average filter. These averages were then used as the temporal features (27). This process led to a sampling frequency of 50.8 Hz, and hence we call it dataset *f50_avg*. This method is justified based on the knowledge that the P300 response - the signal of interest - is of low frequency relative to the sampling frequency (21, 27). Each trial, comprising 60 stimuli flashes, was then segmented into 60 segments from 100 milliseconds pre-stimulus to 700 milliseconds post-stimulus. Each segment represents an input example, *x*, whose corresponding output, *y*, is either 1 if the participant attended this location or 0 otherwise. Each segment was then baseline-corrected by subtracting the average of the pre-stimulus interval. Since the dataset was imbalanced, we oversampled the minority class in the training set by randomly replicating examples until the fractions of both classes were equal before training the neural network models (18). The input data matrix was then normalized by subtracting the mean and dividing by the standard deviation, both determined from the training set.

### CONVOLUTIONAL NEURAL NETWORKS (CNNS)

A convolutional layer of a CNN is formed from multiple feature maps. Each feature map comprises artificial neurons that share the same set of weights, but they apply their weights to a different patch of the input. These weights are called a filter, a kernel, or a feature detector, and their input patches are the equivalent of receptive fields, in the analogy between CNNs and the visual system. Each feature map thus represents the result of a feature detector striding through the input to scan it and outputs a similarity measure to this feature in all possible locations. These filters are learned from the data with backpropagation and gradient descent in the same way as fully connected networks (11). That is why a CNN is considered a feature extractor as well as a classifier or regressor, based on the task for which it is used. Convolutional layers are usually followed by pooling layers. A pooling layer is a downsampling operation that is applied to each feature map. The pooling operation is defined by a pooling size and a pooling stride. The pooling size controls the input patch size, and the pooling stride controls how to move to the next input patch. The pooling layer serves to reduce the dimensionality of the network and therefore the complexity of the model. It also causes the model to be spatially invariant.

#### The branched model

We designed a model consisting of five convolutional layers plus the input and output layers (Fig. 1). The first convolutional layer is purely spatial, i.e., the filter extends through all the channels at each time point separately. On top of the output of the spatial filters layer, we implemented two parallel temporal convolutional layers: one with a small filter size that is equal to 100 milliseconds and the other one with a large filter size that is equal to roughly 400 milliseconds. EEG data are thought to be formed from overlapping harmonics of different frequencies. The small and large filters should be able to detect high frequency and low frequency patterns in the data, respectively (48). We then added two more deep convolutional layers after the small filters layer, so the network is deep and wide at the same time. We used *tanh* as the non-linear activation function for each layer, except for the output layer, for which we used a sigmoid function. The reason behind choosing *tanh* as an activation function is that its output range includes positive and negative values with a mean of zero. This is more suitable for representing features of EEG data, which include positive and negative values, than the commonly used *ReLU* function (16), whose output range has only positive values. Additionally, we used batch normalization (22) and dropout (46) of 0.5 after each layer to accelerate training and regularize the network. Moreover, we used L2 regularization with regularization strength 0.01 for each layer in the network.

**Fig. 1.**
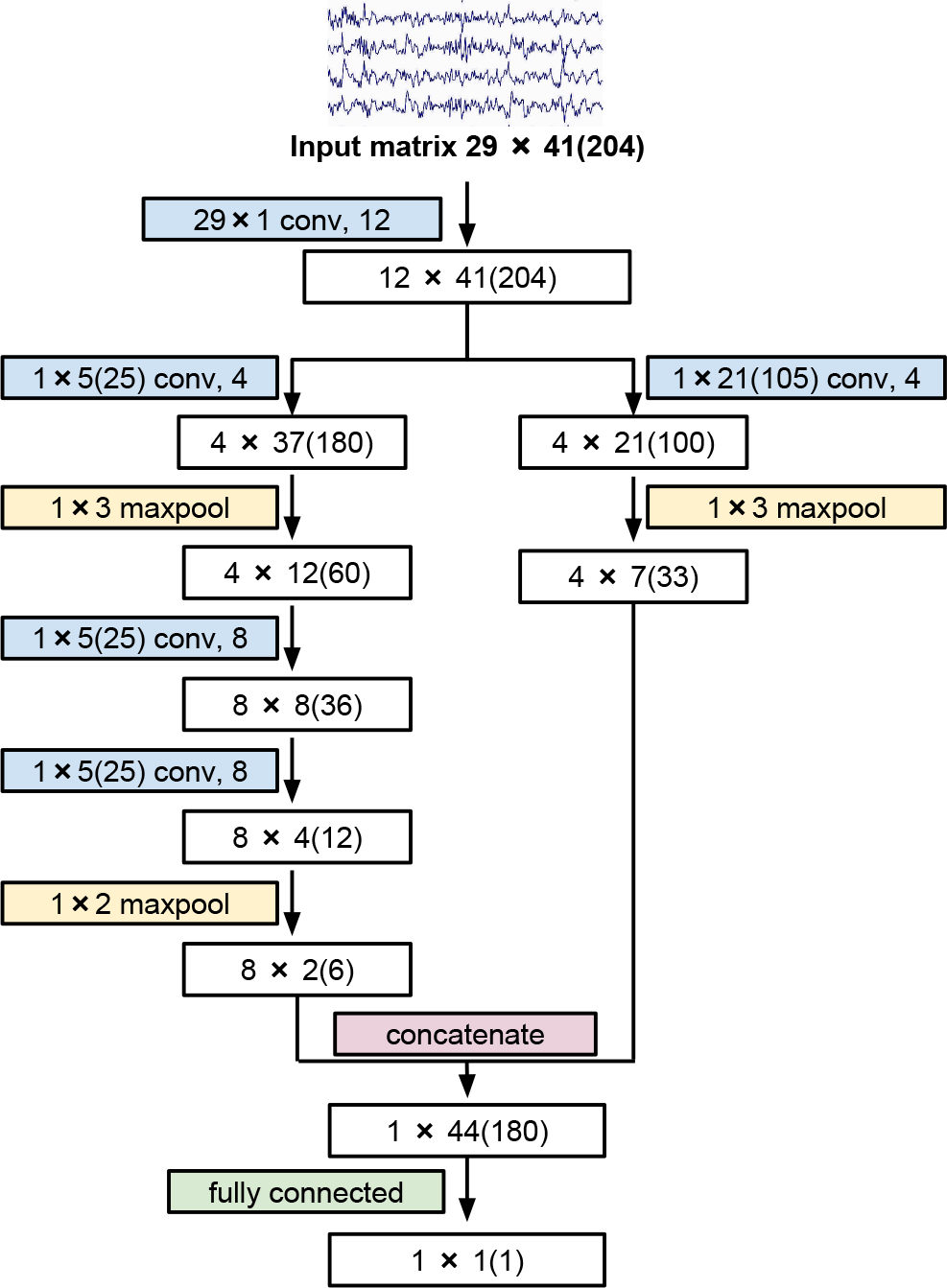
A diagram of the main layers of the branched model. White boxes display the output sizes of the corresponding layers (in colored boxes) for the lower- and higher-dimensional (in parentheses) datasets. Every convolutional layer (in blue) is followed by batch normalization, *tanh* activation and dropout layers.

We validated the impact of different model components by omitting single components, retraining the model and comparing the results. We therefore trained 5 additional models:

- **No_BN**: batch normalization was omitted.
- **No_Dropout**: dropout was omitted.
- **Not_Branched**: the branch with the large temporal filters was omitted to validate the impact of the parallel convolutions.
- **Not_Deep**: the last two layers after the small temporal filters branch were omitted so that the model was formed only from the two parallel convolutional layers on top of the spatial convolutional layer.
- **ReLU**: we replaced the *tanh* activation function with the *ReLU* function (16).
- **ELU**: we replaced the *tanh* activation function with the *ELU* function (10).

We compared the performances of the branched model with a general CNN model for EEG-based BCIs, the EEGNet model (28), in order to test the potential of CNNs in general, and also with a traditional machine learning technique frequently found in the BCI literature, linear discriminant analysis (LDA) (23) and LDA with shrinkage (4).

### CROSS-VALIDATION SCHEMES

To evaluate the performance of the models, we validated within-subject models as well as cross-subject models. For each participant, a model was trained only using the participant’s individual data, rendering the model subject-specific. Since the data are limited per participant, we implemented a 5-fold sequential cross-validation scheme. For LDA, five models were trained per participant, each trained with data from four folds as the training set, and the models were then tested on the data of the held-out fold. For CNN models, we further divided the data from the four folds in a stratified random manner into 80% training data and 20% validation data.

To investigate the model’s transfer learning ability, we implemented a leave-one-subject-out cross-validation scheme in which one participant’s data were held back as a test set, and the data from the remaining 18 participants were used as training and validation sets. We then fine-tuned each cross-subject model with a subset from the test participant’s data in order to evaluate whether we could achieve better performance than with random initializations.

### SALIENCY MAPS

We calculated a within-subject saliency map for each participant based on the participant’s own data and one cross-subject saliency map based on the data of all the participants. In order to calculate the within-subject saliency maps, we trained one model per participant by dividing the participant’s data in a random stratified manner into a 90% training set and a 10% validation set. For the cross-subject saliency map, we trained one cross-subject classifier by also randomly dividing the data of all the participants in a random stratified manner into a 90% training set and a 10% validation set. The saliency maps were then calculated by averaging over all the saliency maps of the target examples in the dataset.

### IMPLEMENTATIONS AND TRAINING SETTINGS

LDA models were implemented using the Scikit-learn library (36). CNN models were implemented using Keras Api (9) with Tensorflow backend (1). The training experiments were conducted on our university’s Medusa CPU cluster.

To optimize the models, we used the Adam optimizer with the recommended settings (learning rate = 10^−3^, *β*_1_ = 0.9, *β*_2_ = 0.999 and *ϵ* = 10^−8^) (24). For fine-tuning, we used stochastic gradient descent with learning rate = 10^−4^ and momentum = 0.9. We used a decaying learning rate scheme that halves the learning rate if there was no decrease in the validation loss for 10 epochs. The early stopping technique was applied to avoid overfitting and to reduce the computational time by stopping the training when the validation loss did not decrease for 20 epochs. The batch size for the mini-batch gradient descent was set to 64 examples. The weights of the networks were initialized with the Glorot Uniform method (15).

## RESULTS

### MODEL PERFORMANCE

With our branched model, we achieved average decoding accuracies of 90.6% (SD: 10%) and 90.7% (SD: 8.6%) on the lower-dimensional and higher-dimensional datasets, respectively (Fig. 2a). The diagrams show the decoding accuracies averaged over the participants. Decoding accuracy indicates the prediction of the target stimulus location as obtained from the binary classification of each stimulus, i.e., the chance level is at 8.3%. Since each stimulus location was flashed five times, we averaged the model outputs of the five repetitions to obtain more robust and accurate predictions. Asterisks above the bars represent the p-value of a Wilcoxon signed-rank test between this model and the branched model Bonferroni corrected for multiple comparisons (12). The branched model yielded significantly higher decoding accuracies than the EEGNet model on the *f50_avg* dataset and than all three models on the *f250* dataset. It is important to note that the branched model was designed for the *f50_avg* dataset and then simply scaled up to fit the bigger *f250* dataset. EEGNet was originally designed for a dataset with a sampling frequency of 128 Hz (28), and therefore, we scaled it up to fit the *f250* dataset and down to fit the *f50_avg* dataset. The EEGNet model achieved significantly better decoding accuracy (p < 0.001) on the *f250* dataset than on the *f50_avg* dataset. In contrast, the difference of decoding accuracies between lower- and higher-dimensional datasets was not significant with the branched model. For the LDA classifier, the shrinkage regularization improved the decoding accuracy (p<0.001) on both datasets, which is particularly clear on the *f250* dataset with higher dimensions, as expected. Nevertheless, on the higher-dimensional dataset, CNNs performed significantly better than LDA with shrinkage.

**Fig. 2.**
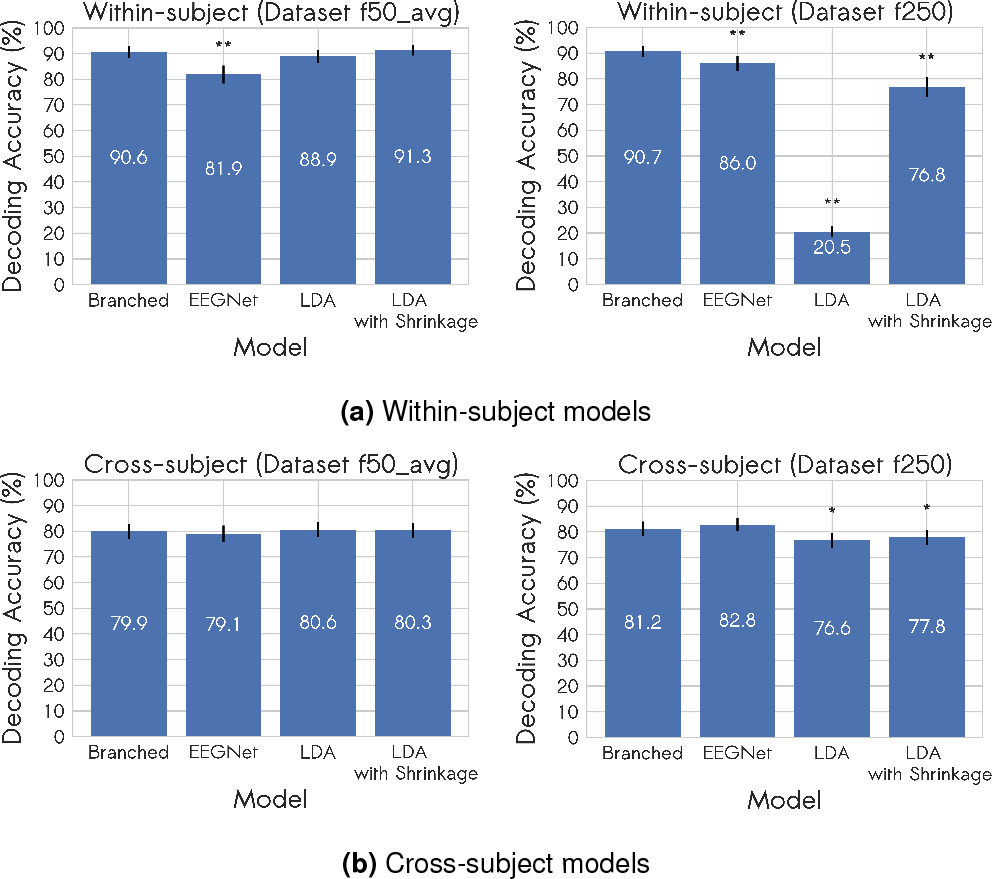
The decoding accuracies of within-subject (a) and cross-subject (b) models applied to the *f_50avg* (left) and *f250* (right) datasets and averaged over the 19 participants. Error bars indicate standard error of the mean. Asterisks denote the statistical significance of the difference between this particular bar and the first bar representing the branched model. (*: p < 0.05, **: p < 0.001 corrected for multiple comparisons)

With the cross-subject branched model, we achieved decoding accuracies of 81.2% (SD: 12.4%), and 79.9% (SD: 12.1%) using the *f250* and *f50_avg* datasets, respectively (Fig. 2b). There was no significant difference between the branched model and any of the other three models on the *f50_avg* dataset, but it still yielded significantly higher accuracy than both LDA classifiers on the *f250* dataset. While the branched model only showed a tendency towards better performance on the high-dimensional *f250* dataset, the EEGNet model achieved a significantly higher accuracy (p < 0.05).

Moreover, the effect of shrinkage on the LDA classifier is not significant, as in the case of within-subject classifiers on both datasets. This is because shrinkage works in situations where the number of training examples is small compared with the number of features. In this case, when the data were pooled from 18 participants, this ratio was high and therefore, LDA was less prone to overfitting, and the regularization was not advantageous.

### IMPACT OF MODEL COMPONENTS

We investigated the impact of different components of the branched model by comparing the decoding accuracies for different alternative within-subject models, in which we omitted or changed one of the model components (Fig. 3a). Again, we used Wilcoxon signed-rank tests with Bonferroni correction to test the significance of the differences. Omitting the dropout and changing the activation function to *ReLU* caused the most significant negative effect on both datasets. Removing the large temporal filters branch induced a significant negative effect on the *f50_avg* dataset. The effect of batch normalization was mainly negligible and differed across the datasets. Using the *ELU* activation function led to a non-significant improvement in the performance across both datasets. The same effect was observed when removing the deep convolutional layers (*Not_deep*).

**Fig. 3.**
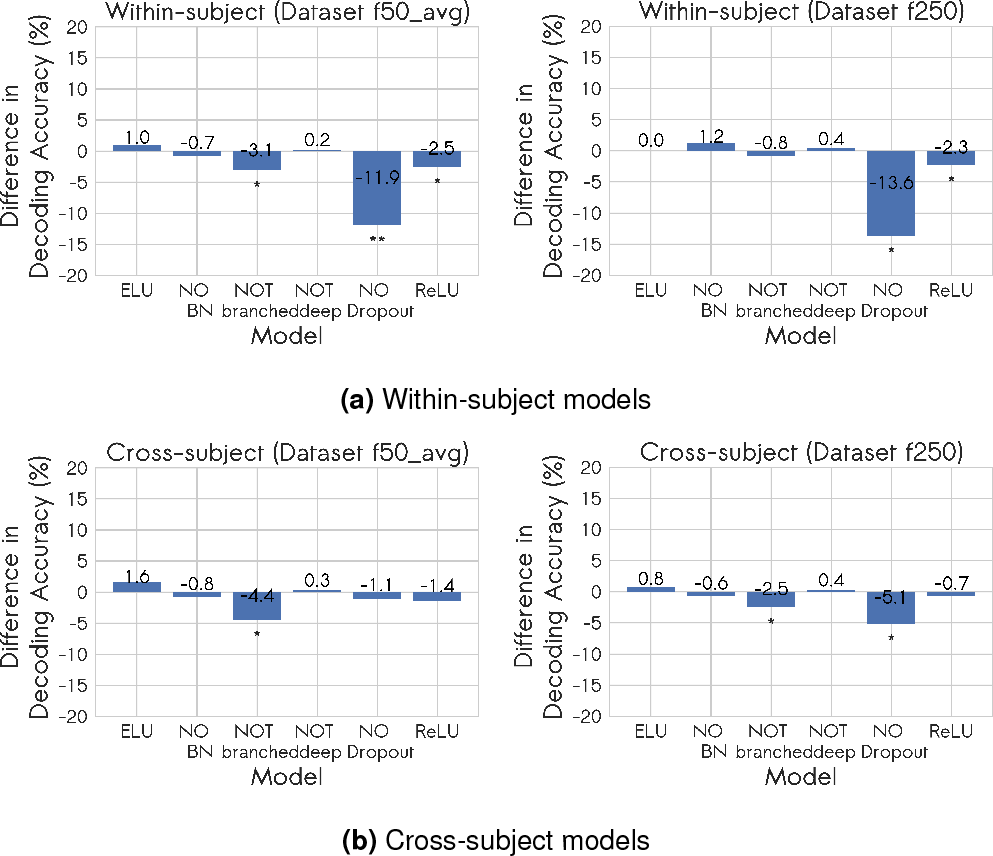
Effect of different model components on the decoding accuracies of within-subject (a) and cross-subject (b) models applied to *f_50avg* (left) and *f250* (right) datasets and averaged over the 19 participants. Asterisks denote the statistical significance of the difference between this particular bar and the original branched model configuration (Fig. 1). (*: p < 0.05, **: p < 0.001 corrected for multiple comparisons)

We also tested the impact of the model components for the cross-subject models. We observed a similar trend to that seen with the within-subject models: mainly the performance decrease of the *ReLU* activation function, *No_dropout*, and *Not_branched* and the performance increase of *Not_deep* and the *ELU* activation function (Fig. 3b).

### FINE-TUNING AND TRANSFER LEARNING

Training cross-subject branched models yielded average decoding accuracies of 79.9% and 81.2% on the *f50_avg* and *f250* datasets, respectively (Fig. 2b). Starting with these models, we continued to fine-tune them with the test participants’ data. We used 10, 20, 30, 50, 60 and 70 trials to further train the models and re-evaluated them using the remaining data (Fig. 4). This approach simulates the situation when a new participant uses the model out-of-the-box and tunes it with subject-specific training trials. Incrementally adding subject-specific trials, a significant improvement in accuracy was observed for both datasets from 30 additional trials. After using 70 trials, the models achieved decoding accuracies of 91.8% (SD: 9.2%), and 91.1% (SD: 9.6%) on the *f50_avg* and *f250* datasets, respectively.

**Fig. 4.**
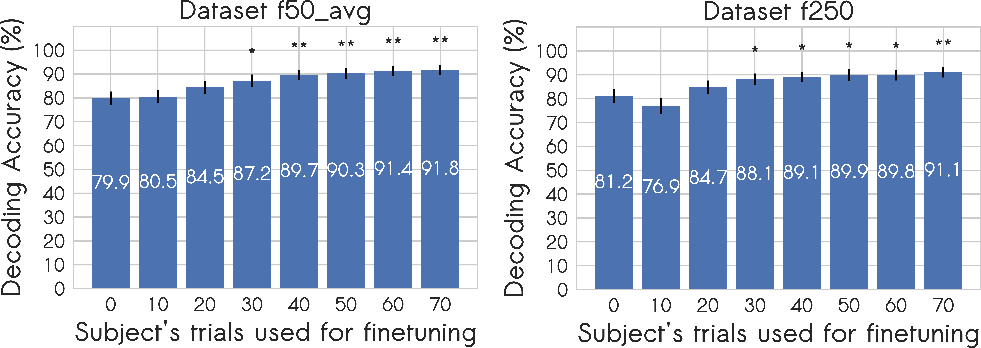
Effect of fine-tuning the cross-subject branched model with the test participant’s own trials. Asterisks denote the statistical significance of the difference between this particular bar and the original cross-subject branched model (0 trials). (*: p < 0.05, **: p < 0.001 corrected for multiple comparisons)

These results cannot be compared directly to the results of the cross-validation scheme of the within-subject models trained from random initializations because of the difference in the test schemes. We therefore trained within-subject models with the same cross-validation scheme but starting from the cross-subject model as a starting point instead of random initializations. These new within-subject models achieved decoding accuracies of 91.9% (SD: 8.5%), and 93% (SD: 6.7%) on the *f50_avg* and *f250* datasets, respectively. They showed improvements over the 90.6% and 90.7% reported for cross-subject models (Fig. 2b). These improvements were significant for both datasets (*f50_avg*: p < 0.05; *f250*: p < 0.001).

### SALIENCY MAPS

To create the within-subject saliency maps, the single-example saliency maps (see Fig. 5) for all the target examples in the participant’s dataset were averaged. We examined the within-subject saliency maps of two participants for the branched model, who had decoding accuracies of 98.4% and 73.6% respectively, and were chosen to illustrate saliency maps for high- and low-performing participants (Fig. 6). The features expected to drive the classification using this paradigm are identifiable on inspection of the saliency map of the high-performing participant: central and parietal electrode locations (FC2, CP1, CP2, and P7) show salient features 300-500 milliseconds post-stimulus. The saliency map of the low-performing participant, on the other hand, has a more scattered appearance across electrodes and over time, which could be due to noisy data leading to low decoding accuracy and poor noisy saliency maps.

**Fig. 5.**
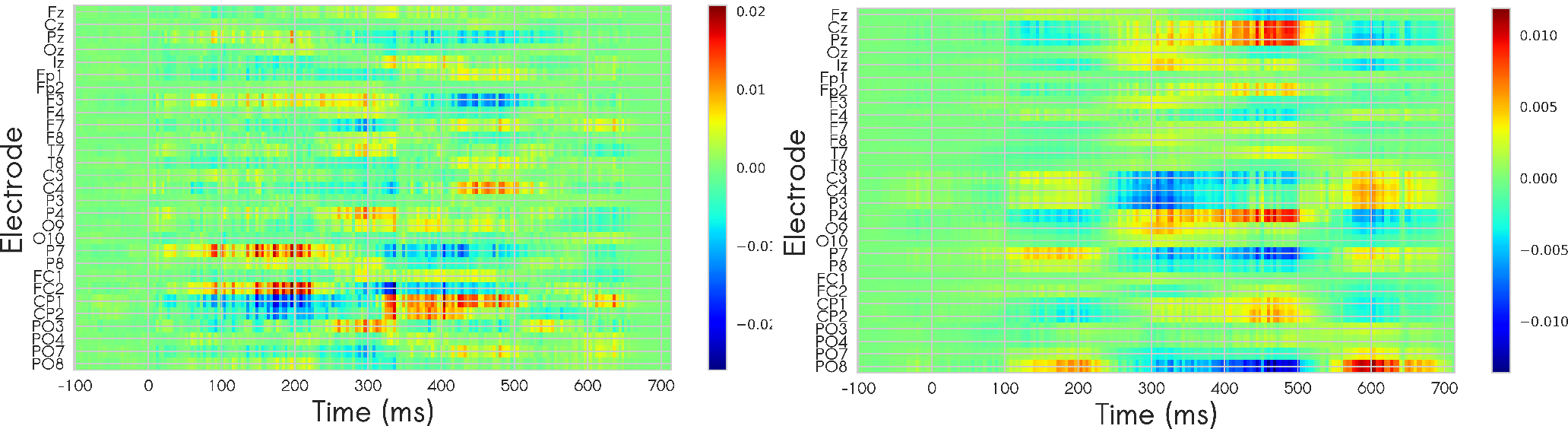
Single-example saliency maps, calculated for the same target input example. Left: generated for a within-subject branched model. Right: generated for a cross-subject branched model. The maps show the positive gradients in red and the negative gradients in blue. Positive and negative gradients indicate the direction in which we would have to change this feature to increase the conditional probability of the attended class given this input example and their magnitude shows the size of this effect.

**Fig. 6.**
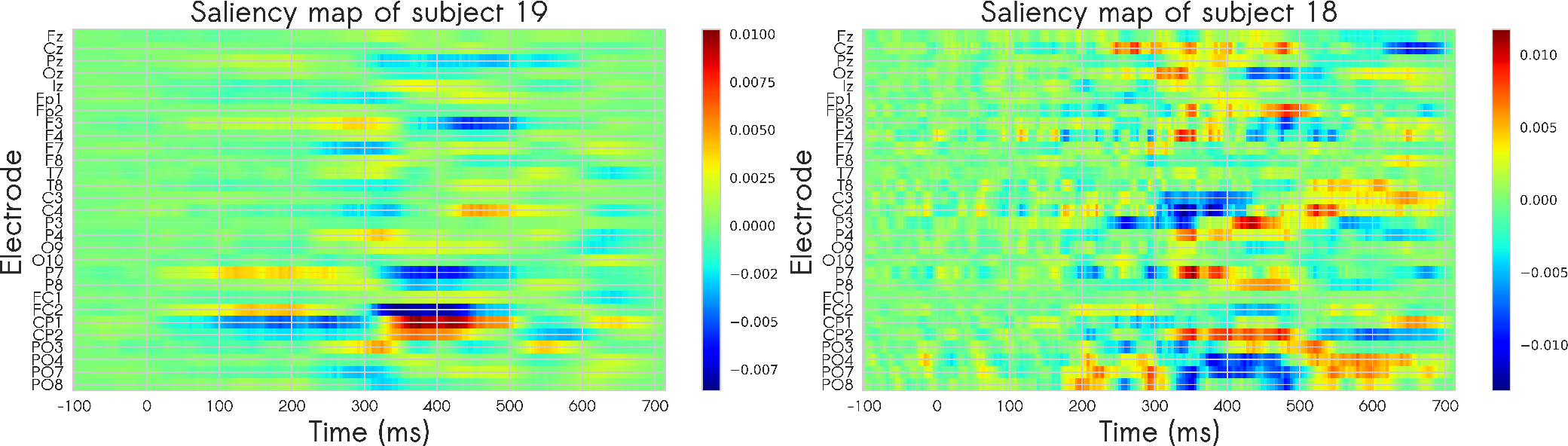
Within-subject saliency maps of the branched model for two different participants, generated by averaging over all single-example saliency maps of target examples for each participant. The participant on the left achieved high decoding accuracy (98.4%) and the participant on the right achieved low decoding accuracy (73.6%).

The cross-subject saliency map (Fig. 7) is more suitable for visualizing more general task-relevant features, as the cross-subject model is less likely to pick up subject-specific, task-irrelevant features (see the difference for a single input example on Fig. 5). These subject-specific features could help improve the decoding accuracy of within-subject models but at the expense of their generalizability to other participants. Examining the grand average ERP at the most discriminative electrode, we observed some resemblance between the ERP waves and the gradients shown on the saliency map (Fig. 7). It is important to note that the resemblance is not exact as the models were trained on a set of single trials which differed from the average wave. We observe that the grand ERP for the target stimuli shows higher amplitude than the grand ERP of the standard stimuli around 200 milliseconds and after 300 milliseconds and lower amplitude around 100 milliseconds and 300 ms. These differences are reflected on the saliency maps by positive and negative gradients. Note that the negative difference at around 100 milliseconds is not represented on the saliency map which could be because the difference is small and not relevant for the classification task.

**Fig. 7.**
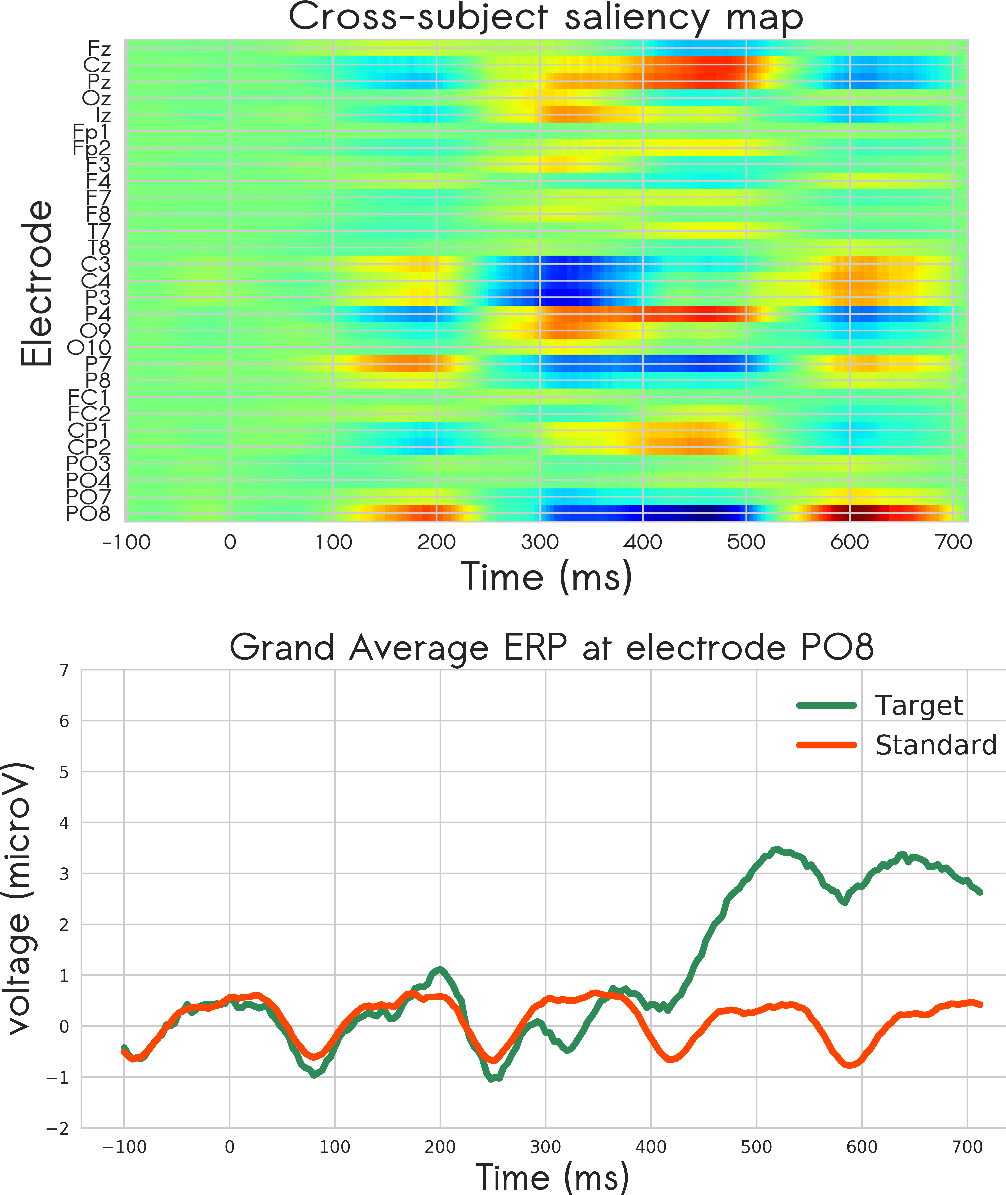
Cross-subject saliency map of the branched model (top). The lower panel shows the grand ERPs recorded from the most discriminant electrode as determined by the saliency map.

### SPATIAL AND TEMPORAL FEATURES EXTRACTION

CNN models were found to be especially powerful in dealing with high-dimensional data (*f250*), as they are capable of automatically extracting task-relevant features from the data. Therefore, besides gaining insights from the saliency maps by visual inspection, it would be also useful to quantify them in order to learn the relevant spatial and temporal features that drive the network decision. To this end, we quantified the saliency maps by computing the average of their absolute values across space and time.

Spatial features are defined as the most discriminative electrodes across the whole trial time period. The eight most discriminative electrodes for each participant separately computed on the within-subject saliency maps are illustrated in Fig. 8a. We can observe that they were variable across participants, but were mainly located around central and parietal electrodes, with some participants having temporal, occipital, or frontal extensions. The eight most discriminative electrodes extracted from the cross-subject saliency map also showed central and parietal distribution (Fig. 8b).

**Fig. 8.**
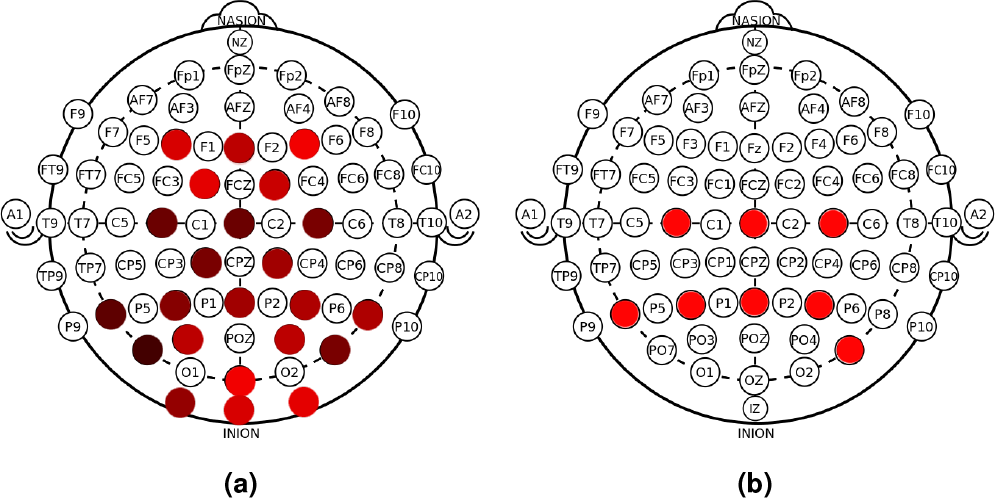
(a) The eight most discriminative electrode positions for within-subject models. The darker the electrode, the more frequent it appeared in the within-subject eight most discriminative electrodes. (b) The eight most discriminative electrode positions for the cross-subject model.

Temporal features are defined as the most discriminative time periods for the detection of the target stimuli. The most discriminative 100 milliseconds time window across all the electrodes for within-subject models and for the cross-subject model fell in the interval between around 300 and 500 milliseconds post-stimulus (Fig. 9).

**Fig. 9.**
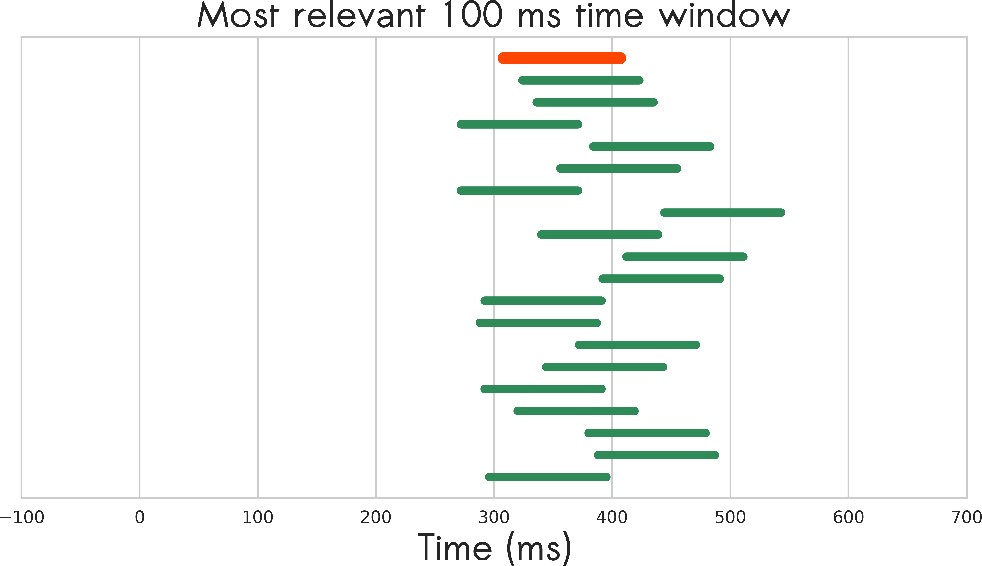
Each green line shows the most discriminative 100 milliseconds time window for one within-subject model. The orange line shows the most discriminative 100 milliseconds for the cross-subject model.

These spatial and temporal features correspond with the well-established spatial distribution and time period of the P300 component (38). The distribution of the normalized gradients, averaged across all the electrodes for each time point for both the within-subject and cross-subject saliency maps across time, resembled a trimodal distribution, with the highest contribution coming from the time period between 300 and 500 milliseconds, followed by contributions from the time period between 100 and 200 milliseconds and around 600 milliseconds (Fig. 10). Each peak in the distribution could be interpreted as reflecting a specific neural process in the brain that is modulated by covert attention at a specific time period.

**Fig. 10.**
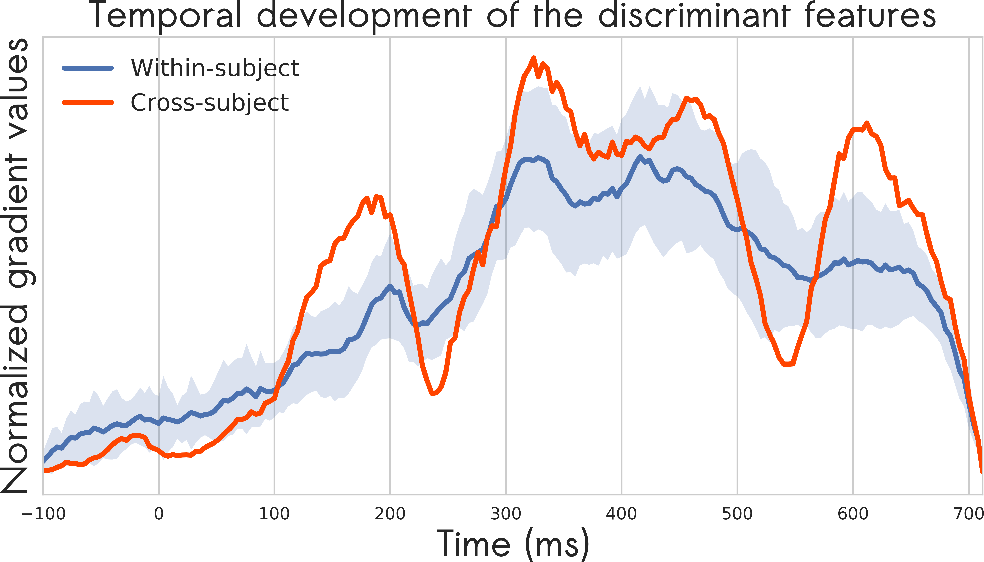
Each time point represents the normalized average of the absolute value of the gradients across space. It shows how discriminative each time point is and therefore how the discriminability of the EEG features develops over time. The within-subject curve is averaged across the 19 participants with the shade denotes the standard deviation.

### VALIDATION OF THE EXTRACTED FEATURES

We validated the interpretation of the absolute values of the gradients in the saliency maps as the discriminative power of the features by comparing the decoding accuracy using all electrodes with that obtained using only the eight most and the eight least discriminative electrodes, setting all other electrodes to zero in the latter cases (Fig. 11a). Inside the cross-validation scheme of evaluating the performance of the within-subject branched models, we used the trained models to generate saliency maps, from which we computed the rank of the electrodes (as mentioned previously and shown in Fig. 8a). Visual inspection clearly reveals a difference in the decoding accuracies in the case of using the eight most discriminative versus the eight least discriminative electrodes (Fig. 11b). Using only 8 out of the 29 electrodes means using only 27.5% of the features. Using the eight most discriminative electrodes, we retained 72% of the decoding accuracy versus only 16% in the case of using the eight least discriminative electrodes.

**Fig. 11.**
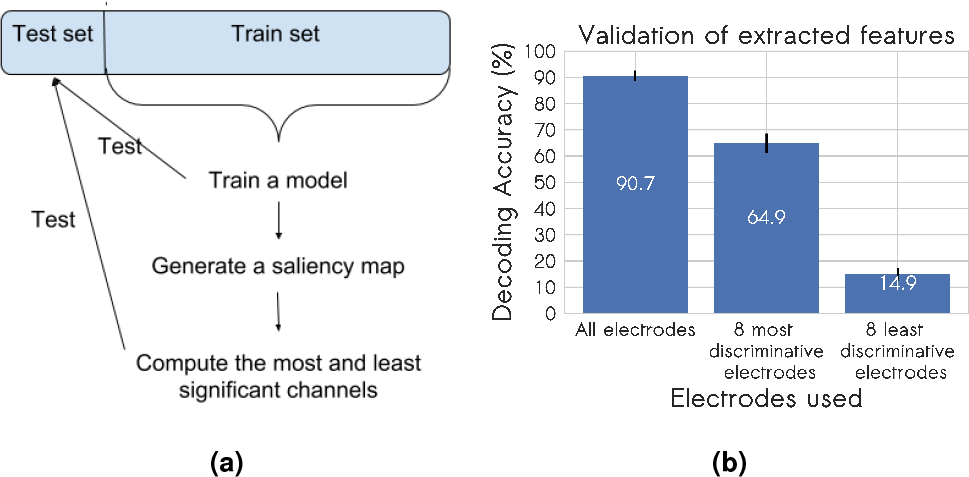
Panel (a) shows the scheme used for validation of the discriminative power of the extracted features. Plot (b) compares the decoding accuracies of the full set of electrodes, the eight most discriminative electrode sites and the eight least discriminative electrode sites.

Furthermore, we compared the eight most discriminative electrodes determined with the saliency map approach to an approach that is based on considering the weights (spatial filters) of the first hidden layer (8, 31). We trained within-subject LDA classifiers with shrinkage using only the eight most discriminative electrodes determined by both approaches. There was no significant difference between the two approaches, as using saliency map and spatial filter approaches yielded decoding accuracies of 80.70% and 81.19%, respectively (*p* = 0.6). Moreover, by comparing the eight most discriminative electrode sites for the cross-subject model, there was only a difference in one electrode (replacing electrode C3 from Fig. 8b by electrode CP2).

## DISCUSSION

We have demonstrated that CNNs can be employed to decode electrophysiological signals for P300 BCI applications and can furthermore provide insights into the underlying brain processing when combined with visualisation using saliency maps. CNNs performed at least on a par with commonly used decoding methods, and their performance was significantly superior when applied to high-dimensional, minimally-processed data on both within-subject and cross-subject decoding tasks. CNNs also achieved higher accuracies on the high-dimensional data than on the low-dimensional data with hypothesis-driven feature extraction. This superior performance can be attributed to their ability to extract optimal relevant features automatically from the data that leverage separability of the data. This applicability to high-dimensional data has particular potential in novel decoding problems, in which we have no a priori information regarding the most discriminative signal. For example, in the covert attention-decoding problem, the use of downsampling after applying a moving average filter is permissible, because it is known that the P300 signal is of low frequency in relation to the EEG acquisition sampling frequency (21, 27). For cases in which the discriminative signals are represented in high frequency bands, this method would cause attenuation of the discriminative information and therefore reduction of the decoding accuracy. CNNs also allowed transfer learning through the process of network fine-tuning. Fine-tuning the cross-subject branched model led to a significant increase in the decoding accuracy of the within-subject models in comparison to training from random initialization. Such transfer learning approach would potentially reduce the time needed for acquisition of training data, which in turn would make BCI systems more accessible for patients who need it.

A limitation of the approach is that optimizing CNN models is a time-consuming and computationally expensive process in comparison to training a simple linear classifier like the LDA. The models usually contain tens or hundreds of hyper-parameters that need to be tuned to achieve reasonable performance. These hyper-parameters are data-dependent and are tuned mainly through intensive initial parameter adjustment, intuition, and experience with the data. There is, however, a growing research interest in automated neural architecture search (31, 37), but it is still computationally expensive.

Furthermore, the results showed that not all the standard choices of these hyperparameters led to the best results. For example, applying the commonly used *ReLU* activation function led consistently to poorer results in comparison to *tanh* or *ELU* activation functions. This is probably because the *ELU* and *tanh* activation functions have similar properties, such that they both have positive and negative output values in contrast to the *ReLU* activation function, which has only positive output values. Using the dropout regularization technique, however, caused significant improvement of the models performance, which was particularity observable when the dataset was small (within-subject) and high-dimensional (*f250*). Moreover, using the parallel convolutional layers, which were motivated by the nature of the EEG data, led to more significant improvement than using deep sequential layers. This suggests that parallel convolutions are more suitable for EEG data decoding than the deep ones. Additionally, batch normalization did not provide a significant improvement in contrast to what was reported in (31), which fits with the notion that designing the best network architecture is data-dependent.

The fact that CNNs can work directly on raw, high-dimensional data and automatically extract relevant features poses an opportunity to discover information about the cognitive task at hand, if we can visualize these relevant features. We showed how saliency maps can be used for feature visualization in EEG data. It was possible to visualize the discriminative power of the features on a single example level, which is suitable for single-trial EEG analysis, on a within-subject level, which is relevant for studying between-subject variability, and on a cross-subject level, which is necessary for study ing general task-relevant features. Moreover, quantifying the discriminability of the features across electrodes or time provided insight into the task-relevant electrodes and time periods. The cross-subject saliency map showed that the eight most discriminative electrodes were PO8, P4, P7, Pz, Cz, C3, P3 and C4 (Fig. 8b). This finding is in agreement with the 8-channel montage optimized by (27) and (26) for P300 spellers in 5 electrodes (PO8, Cz, Pz, P3, and P4). Additionally, their montage included Oz, PO7, and Fz, which with our approach were ranked at positions 18, 13, and 16 respectively. The saliency map approach determined, on the other hand, the electrodes P7, C3, and C4 to be more predictive, in agreement with the findings of Brunner et al. (6), who reported that this 8-channel montage is suboptimal when the participants do not fixate their gaze on the stimulus. The branched model identified central and parietal electrodes as being more discriminative than occipital and parieto-occipital ones. These locations are in line with the parietal and central distribution of the P300 component, which is known to be elicited in covert attention paradigms, in contrast to the occipital and parietooccipital distribution of VEPs, known to be modulated in overt attention paradigms (49). The difference between covert (as performed here) and overt attention paradigms is that participants foveate the target stimulus when attention is overt. Furthermore, quantifying the cross-subject saliency map across time (Fig. 9 and Fig. 10) showed that the most discriminative time period started from 300 milliseconds post-stimulus. There are still contributions from the time period between 100 and 200 milliseconds poststimulus, which corresponds to the attentional modulations of VEPs, but they are less prominent. It has been reported that for overt attention tasks, the time period from 180 to 250 milliseconds post-stimulus is the most informative, when P2 (second positive component) and N1 (first negative component) occur. On the contrary, for covert attention tasks the post-300 milliseconds time period is the most informative, in which the P300 component occurs (49).

Within-subject saliency maps could also be used to investigate the variability between participants. Examination of the variability of the eight most discriminative electrodes across participants showed that most participants have a central and parietal distribution of the relevant electrodes with occipital extension and sometimes a frontal one (Fig. 8a). The discriminative time periods are mostly concentrated in the time period between 300 and 500 milliseconds post-stimulus across participants (Fig. 9, Fig. 10). Determining between-subject variability and within-subject variability (by comparing the saliency maps of early and late trials) could have a potential application in the development of biomarkers to diagnose and monitor the prognosis of certain neurological or psychiatric disorders, e.g., prolonged P300 latency in dementia patients (38).

Another approach for determining the most discriminative electrodes is to consider the weights of the first hidden layer (8, 31) and interpret them as spatial filters. Taking the average of the absolute values of the layer’s spatial filters gives an estimate of the discriminative power of the electrodes (8). This approach, however, has disadvantages. First, it is architecture-dependent, because it can only be applied to the first hidden layer in cases in which this layer is a spatial convolution layer. Secondly, it does not account for the transformation that will be applied to the output of this layer in the subsequent layers. Thirdly, it can only extract spatial features but not temporal or spatiotemporal features, so for example, it can not provide information about the most discriminative time periods. Comparing the saliency map approach for relevant feature extraction with this approach showed that the saliency map approach is at least as effective in extracting spatial features. Additionally, it has the advantages of being architecture independent and able to extract temporal features as well, which can potentially help explore the brain dynamics associated with a cognitive task.

It is also worth noting that these saliency maps can potentially offer an objective heuristic for doing ERP research. An objective approach could be achieved through training a CNN model to classify the different conditions under investigation, and by visualizing the saliency maps for each class, we could gain an indication of where and when to look for different neural correlates for the conditions in a data-driven way.

## Conclusion

Convolutional neural networks performed significantly better than traditional decoding techniques on the high-dimensional data. However, this superior performance comes at the expense of time and computational costs. Using the standard recommended model components does not guarantee the best performance and therefore manual optimization is still needed. More studies that validate and potentially replicate our design recommendations are also still needed. Thus, we believe that for cases, in which we have domain knowledge of the classification problem, like in the case of P300 classification, using a simple linear classifier on top of a feature extraction method that is based on this domain knowledge, offers a more practical solution. On the other hand, in novel cases, in which we do not have sufficient prior knowledge of the task-relevant effects on the EEG signal, using CNN models is justifiable and recommended. Training CNN models in these cases is not only more likely to produce performances competitive with other methods, but also to provide insight into the task-relevant features through visualization techniques like saliency maps.

## Future research

For such models to be applicable in real world situations, they need to be tested on other datasets. Further investigation is required to establish whether they could be used ‘out of the box’ with reasonable performance, needing only some fine-tuning with a small amount of data, or whether retraining from scratch is necessary.

Moreover, there is still room for improvement, through exploring different neural network architectures, such as recurrent neural networks (RNN) (17), long short term memory networks (LSTM) (19), hybrid networks, and automatic neural architecture search techniques (31, 37).

Using CNN models for EEG decoding still needs further development before such models could become the standard approach and replace other techniques. The research in this area needs to go beyond intensive initial parameter adjustment and start to investigate the underpinnings of the models when they are applied to EEG data.

## References

1. Abadi, M., Barham, P., Chen, J., Chen, Z., Davis, A., Dean, J., Devin, M., Ghemawat, S., Irving, G., Isard, M., Kudlur, M., Levenberg, J., Monga, R., Moore, S., Murray, D. G., Steiner, B., Tucker, P., Vasudevan, V., Warden, P., Wicke, M., Yu, Y., and Zheng, X. (2016). Tensorflow: A system for large-scale machine learning. In 12th USENIX Symposium on Operating Systems Design and Implementation (OSDI 16), pages 265–283, Savannah GA. USENIX Association.

2. Amini, Z., Abootalebi, V., and Sadeghi, M. T. (2010). A comparative study of feature extraction methods in P300 detection. 2010 17th Iranian Conference of Biomedical Engineering ICBME, (November):1–4.

3. Ancona, M., Ceolini, E.,Öztireli, C., and Gross, M. (2018). Towards Better Understanding of Gradient-Based Attribution Methods. In International Conference on Learning Representations, number Section 3, pages 1–16.

4. Blankertz, B., Lemm, S., Treder, M., Haufe, S., and Müller, K. R. (2011). Single-trial analysis and classification of ERP components - A tutorial. NeuroImage, 56(2):814–825.

5. Blankertz, B., Muller, K., Krusienski, D., Schalk, G., Wolpaw, J., Schlogl, A., Pfurtscheller, G., Millan, J., Schroder, M., and Birbaumer, N. (2006). The BCI Competition III: Validating Alternative Approaches to Actual BCI Problems. IEEE Transactions on Neural Systems and Rehabilitation Engineering, 14(2):153–159.

6. Brunner, P., Joshi, S., Briskin, S., Wolpaw, J. R., Bischof, H., and Schalk, G. (2010). Does the ‘P300’ speller depend on eye gaze? Journal of Neural Engineering, 7(5).

7. Carabez, E., Sugi, M., Nambu, I., and Wada, Y. (2017). Convolutional Neural Networks with 3D Input for P300 Identification in Auditory Brain-Computer Interfaces. Computational intelligence and neuroscience, 2017:8163949.

8. Cecotti, H. and Graser, A. (2011). Convolutional Neural Networks for P300 Detection with Application to Brain-Computer Interfaces. IEEE Transactions on Pattern Analysis and Machine Intelligence, 33(3):433–445.

9. Chollet, F. et al. (2015). Keras. https://keras.io.

10. Clevert, D.-A., Unterthiner, T., and Hochreiter, S. (2016). Fast and Accurate Deep Network Learning by Exponential Linear Units (ELUs). In International Conference on Learning Representations.

11. Courville, I. G., Bengio, Y., and Aaron (2016). Deep Learning. MIT Press.

12. Demšar, J. (2006). Statistical Comparisons of Classifiers over Multiple Data Sets. Journal of Machine Learning Research, 7(Jan):1–30.

13. Di Russo, F., Martinez, A., and Hillyard, S. A. (2003). Source analysis of event-related cortical activity during visuo-spatial attention. Cerebral Cortex, 13(5):486–499.

14. Farwell, L. and Donchin, E. (1988). Talking off the top of your head: toward prothesis utilizing event-related brain potentials. Electroencephalography and clinical Neurophysiology., 70(May):510–523.

15. Glorot, X. and Bengio, Y. (2010). Understanding the difficulty of training deep feedforward neural networks. In Teh, Y. W. and Titterington, M., editors, Proceedings of the Thirteenth International Conference on Artificial Intelligence and Statistics, volume 9, pages 249–256. PMLR.

16. Glorot, X., Bordes, A., and Bengio, Y. (2011). Deep sparse rectifier neural networks. In Gordon, G., Dunson, D., and Dudík, M., editors, Proceedings of the Fourteenth International Conference on Artificial Intelligence and Statistics, volume 15, pages 315–323. PMLR.

17. Graves, A., rahman Mohamed, A., and Hinton, G. E. (2013). Speech recognition with deep recurrent neural networks. 2013 IEEE International Conference on Acoustics, Speech and Signal Processing, pages 6645–6649.

18. Haibo He, H. and Garcia, E. (2009). Learning from Imbalanced Data. IEEE Transactions on Knowledge and Data Engineering, 21(9):1263–1284.

19. Hochreiter, S. and Schmidhuber, J. (1997). Long short-term memory. Neural computation, 9(8):1735–80.

20. Hubel, D. H. and Wiesel, T. N. (1962). Receptive fields, binocular interaction and functional architecture in the cat’s visual cortex. The Journal of physiology, 160(1):106–54.

21. Intriligator, J. and Polich, J. (1994). On the relationship between background EEG and the P300 event-related potential. Biological Psychology, 37(3):207–218.

22. Ioffe, S. and Szegedy, C. (2015). Batch normalization: Accelerating deep network training by reducing internal covariate shift. In Bach, F. and Blei, D., editors, Proceedings of the 32nd International Conference on Machine Learning, volume 37, pages 448–456. PMLR.

23. James, G., Witten, D., Hastie, T., and Tibshirani, R. (2013). An Introduction to Statistical Learning, volume 103 of Springer Texts in Statistics. Springer New York, New York, NY.

24. Kingma, D. P. and Ba, J. (2015). Adam: A method for stochastic optimization. In International Conference on Learning Representations.

25. Krizhevsky, A., Sutskever, I., and Geoffrey, E. H. (2012). ImageNet Classification with Deep Convolutional Neural Networks. Advances in Neural Information Processing Systems 25 (NIPS2012), pages 1–9.

26. Krusienski, Sellers, Cabestaing, Bayoudh, McFarland, Vaughan, and Wolpaw (2006). A comparison of classification techniques for the P300 Speller. Journal of Neural Engineering, 3(4):299–305.

27. Krusienski, Sellers, McFarland, Vaughan, and Wolpaw (2008). Toward enhanced P300 speller performance. Journal of Neuroscience Methods, 167(1):15–21.

28. Lawhern, V. J., Solon, A. J., Waytowich, N. R., Gordon, S. M., Hung, C. P., and Lance, B. J. (2018). EEGNet: a compact convolutional neural network for EEG-based brain–computer interfaces. Journal of Neural Engineering, 15(5):056013.

29. Lecun, Y., Boser, B., Denker, J. S., Henderson, D., Howard, R. E., Hubbard, W., and Jackel, L. (1990). Handwritten digit recognition with a back-propagation network.

30. LeCun, Y., Yoshua, B., and Geoffrey, H. (2015). Deep learning. Nature, 521(7553):436–444.

31. Liu, H., Simonyan, K., and Yang, Y. (2019). DARTS: Differentiable architecture search. In International Conference on Learning Representations.

32. Luck, S. J. (2005). An Introduction to the Event-Related Potential Technique. Number 3. MIT Press.

33. Luck, S. J. and Kappenman, E. S. (2011). ERP Components and Selective Attention. Oxford University Press.

34. Manyakov, N. V., Chumerin, N., Combaz, A., and Van Hulle, M. M. (2011). Comparison of classification methods for P300 brain-computer interface on disabled subjects. Computational intelligence and neuroscience, 2011:519868.

35. Olah, C., Mordvintsev, A., and Schubert, L. (2017). Feature visualization. Distill. https://distill.pub/2017/feature-visualization.

36. Pedregosa, F., Varoquaux, G., Gramfort, A., Michel, V., Thirion, B., Grisel, O., Blondel, M., Prettenhofer, P., Weiss, R., Dubourg, V., Vanderplas, J., Passos, A., Cournapeau, D., Brucher, M., Perrot, M., and Duchesnay, É. (2011). Scikit-learn: Machine Learning in Python. Journal of Machine Learning Research, 12(Oct):2825–2830.

37. Pham, H., Guan, M., Zoph, B., Le, Q., and Dean, J. (2018). Efficient neural architecture search via parameters sharing. In Dy, J. and Krause, A., editors, Proceedings of the 35th International Conference on Machine Learning, pages 4095–4104. PMLR.

38. Polich, J. (2012). Neuropsychology of P300. The Oxford Handbook of Event-Related Potential Components, 92037:1–67.

39. Qin, Z., Yu, F., Liu, C., and Chen, X. (2018). How convolutional neural network see the world - A survey of convolutional neural network visualization methods. Mathematical Foundations of Computing, 1(2):149–180.

40. Ramadan, R. A. and Vasilakos, A. V. (2017). Brain computer interface: control signals review. Neurocomputing, 223(October 2016):26–44.

41. Reichert, C., Dürschmid, S., Heinze, H.-J., and Hinrichs, H. (2017). A Comparative Study on the Detection of Covert Attention in Event-Related EEG and MEG Signals to Control a BCI. Frontiers in Neuroscience, 11:575.

42. Sainath, T. N., Mohamed, A.-r., Kingsbury, B., and Ramabhadran, B. (2013). Deep convolutional neural networks for LVCSR. In 2013 IEEE International Conference on Acoustics, Speech and Signal Processing, pages 8614–8618. IEEE.

43. Schirrmeister, R. T., Springenberg, J. T., Fiederer, L. D. J., Glasstetter, M., Eggensperger, K., Tangermann, M., Hutter, F., Burgard, W., and Ball, T. (2017). Deep learning with convolutional neural networks for eeg decoding and visualization. Human Brain Mapping.

44. Simonyan, K., Vedaldi, A., and Zisserman, A. (2013). Deep Inside Convolutional Networks: Visualising Image Classification Models and Saliency Maps. In International Conference on Learning Representations.

45. Springenberg, J., Dosovitskiy, A., Brox, T., and Riedmiller, M. (2015). Striving for simplicity: The all convolutional net. In ICLR (workshop track).

46. Srivastava, N., Hinton, G., Krizhevsky, A., Sutskever, I., and Salakhutdinov, R. (2014). Dropout: A Simple Way to Prevent Neural Networks from Overfitting. Journal of Machine Learning Research, 15:1929–1958.

47. Sturm, I., Lapuschkin, S., Samek, W., and Müller, K.-R. (2016). Interpretable deep neural networks for single-trial eeg classification. Journal of Neuroscience Methods, 274:141–145.

48. Supratak, A., Dong, H., Wu, C., and Guo, Y. (2017). DeepSleepNet: A Model for Automatic Sleep Stage Scoring Based on Raw Single-Channel EEG. IEEE Transactions on Neural Systems and Rehabilitation Engineering, 25(11):1998–2008.

49. Treder, M. S. and Blankertz, B. (2010). (C)overt attention and visual speller design in an ERP-based brain-computer interface. Behavioral and Brain Functions, 6:1–13.

50. Uktveris, T. and Jusas, V. (2015). Comparison of Feature Extraction Methods for EEG BCI Classification. Communications in Computer and Information Science, 538:81–92.

51. Vega-Escobar, L., Castro-Ospina, A. E., and Duque-Munoz, L. (2015). Feature extraction schemes for BCI systems. 2015 20th Symposium on Signal Processing, Images and Computer Vision, STSIVA 2015 - Conference Proceedings, (76).

